# Enhanced Workflow for Urinary Extracellular Vesicle Isolation Using Membrane-Sensing Peptides

**DOI:** 10.64898/2026.05.21.726982

**Authors:** Roberto Frigerio, Adele Tanzi, Angelo Musicò, Cristina Grange, Paola Gagni, Vincenza Dolo, Ilaria Giusti, Paolo Arosio, Alessandro Gori, Benedetta Bussolati

## Abstract

Urinary extracellular vesicles (uEVs) represent a promising source of non-invasive biomarkers; however, their clinical translation is still limited by suboptimal isolation strategies, which often suffer from poor scalability, co-isolation of abundant urinary proteins, and bias toward specific EV subpopulations. Here, we employ a membrane-sensing peptide (MSP)-based affinity approach for uEVs isolation, that exploits the highly lipid membranes curvature of EV as universal target, enabling pan-specific capture independent of surface marker expression. MSP-functionalized beads were applied to minimally processed urine samples and benchmarked against differential ultracentrifugation (dUC) and size-exclusion chromatography (SEC). Comprehensive characterization by nanoparticle tracking analysis, transmission electron microscopy, high-sensitivity flow cytometry, single-molecule array (SiMoA), and fluorescence nanoparticle tracking analysis, demonstrated that MSP-based isolation preserves vesicle integrity and maintains the native distribution of canonical tetraspanins (CD9, CD63, CD81), without evidence of subpopulation bias. Notably, MSP-based isolation significantly reduced co-isolated contaminants, such as uromodulin, resulting in improved sample purity. By combining high recovery, improved purity, and operational simplicity, MSP workflow offers practical advantages, including reduced processing time, scalability, and compatibility with standard laboratory equipment, without the need for extensive pre-processing. These properties characterize MSP-based affinity capture as a robust and versatile alternative to conventional uEVs isolation approaches, with strong potential for translational and clinical applications.

## 1. Introduction

Extracellular vesicles (EVs) are membranous nanosized particles that play a crucial role in intercellular communication ^1,2^. Under both physiological and pathological conditions, cells secrete EVs, whose composition mirrors the state and origin of the secreting cells ^3–5^. Owing to their features, EVs have been emerging as non-invasive, wide biomarkers, providing a valuable window into ongoing biological processes^6–10^. Among the bodily fluids harboring EVs, urine has attracted particular interest because of its easy and non-invasive collection, as well as its rich content of biological compounds^11,12^. Indeed, urinary EVs (uEVs) have been shown to carry clinically valuable information reflective of a wide range of pathological conditions, including mainly renal diseases and urogenital tract-related cancers^13^. Moreover, a few recent studies have also explored the potential of uEVss as therapeutic tools, further highlighting their relevance in the field of translational research^14,15^. Despite rapid progress in the field, isolating uEVss remains a major bottleneck^16^.

Ultracentrifugation is the most widely adopted uEVs practice, although its use for clinical translatability suffers from many drawbacks – including labor- and time-intensive protocols, limited scalability in high-throughput biomarker discovery/validation settings, and difficult interlaboratory cross-validation due to different equipment specifications and/or protocol adjustments^17^.

In light of the ongoing standardization efforts by the EV community, an additional challenge concerns the co-isolation of abundant urinary proteins in uEVs preparations obtained by dUC - most notably uromodulin, a highly abundant polymeric urinary protein that can interfere with downstream EV analysis^13^. Likewise, polymer precipitation-based techniques, exploited in many commercial kits and attractive for simplicity and apparent high recovery^18^, resulting in the co-precipitation of proteins and other non-EV particles.

On the other hand, size-exclusion chromatography (SEC) is a preferred EV isolation method to yield EV fractions with limited contamination from soluble proteins^19,20^, but its reliance on pre- and post-concentration steps introduces additional hands-on time, workflow complexity, and reduced uEVs recovery yield – overall questioning its broad application in standardized uEVs workflows^21,22^.

Last but not least, immunoaffinity capture offers several advantages in terms of purity and high-throughput scalability^23,24^, yet it is intrinsically restricted to specific EV subpopulations and, due to cost considerations, is generally limited in input volume processability, while also requiring the absence of competing non-EV soluble markers. As a matter of fact, current limitations in uEVs isolation methods demand improved methodologies to fully unlock the potential of uEVss in translational and clinical applications^11,25–27^.

Herein, we investigate membrane-sensing peptide (MSP)–mediated EV affinity separation as an alternative strategy for uEVs isolation, benchmarking key operational parameters against one of the gold standard practices, i.e., differential ultracentrifugation (dUC). The key rationale for our investigation was that MSP-isolation may combine the best features of other separation methodologies, namely high-volume sample processing and efficient recovery (UC), low protein contamination (SEC), and high-throughput ready operational workflow (immunoaffinity).

Membrane curvature-sensing peptides have recently entered the EV field, yet they are finding increasingly successful applications^28^. Briefly, MSPs leverage the lipid packing defects characteristic of highly curved and tensioned membranes, enabling the insertion of hydrophobic anchors that stabilize EV binding following an initial electrostatically driven interaction. More expanded mechanistic information is reported elsewhere^28,29^.

From a functional perspective, MSPs enable robust capture of EVs independent of their molecular surface profile, acting as “marker-agnostic” affinity probes. Moreover, owing to their scalable and versatile chemical synthesis, they can be seamlessly integrated onto diverse solid supports and are compatible with workflows across a wide range of sample volumes^29–33^.

This study aims to evaluate MSP-based capture as a robust, process-efficient, and scalable strategy for uEVs isolation, addressing key requirements for translational and clinical implementation. To support this,uEVs isolation by MSP-functionalized beads was directly compared with differential ultracentrifugation (dUC) (Fig. 1 and the resulting EV preparations were characterized in terms of size and concentration (NTA, TEM), purity (ELISA quantification of residual uromodulin), and markers profiling (SiMoA, Fluorescence NTA, MACXPlex, and high-sensitivity flow cytometry).

**Figure 1:**
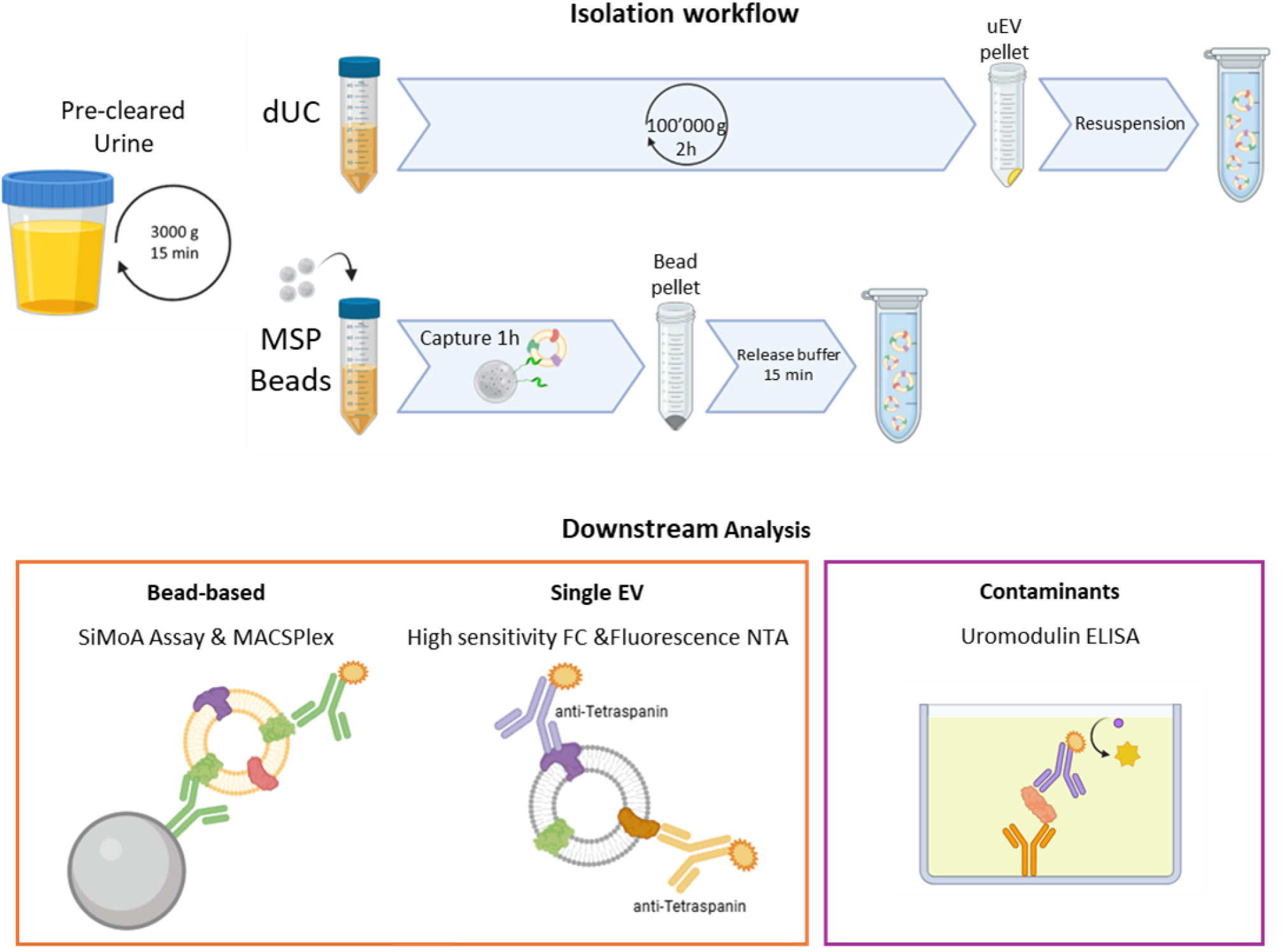
Schematic overview of the comparative workflow for uEVs isolation using dUC and MSP-functionalized beads. The resulting uEVs preparations were analyzed using bead-based assays and single-particle techniques to assess size, concentration, and marker expression, while co-isolated contaminants were quantified by ELISA.

## 2. Materials and Methods

### 2.1 Urine collection and pre-clearing

First morning urines from healthy donors were collected and pre-cleared through centrifugation at 3,000 x g for 15 min, followed by a filtration (0.22 μm pores). Pre-cleared samples were then used for uEVs isolation following different protocols.

### 2.2 uEVs isolation by dUC

Pre-cleared urine samples (20 mL) were ultracentrifuged at 100,000 x g for 2 hours at 4 °C. The resulting pellet was resuspended in 400 μL of reference working buffer (FCS-depleted RPMI 1% DMSO). For comparative assessment, uEVs pellets from different aliquots were resuspended in the release buffer used for EV elution from MSP-beads (RPMI 1% DMSO 200 mM EDTA).

### 2.3 uEVs isolation by SEC

uEVss were isolated using size-exclusion chromatography (SEC) with qEV70 columns (IZON Science), following the manufacturer’s instructions with minor adaptations for urine samples. Briefly, urine was first centrifuged 2,000 × g for 15 min at 4 °C to eliminate cellular debris. The resulting supernatant was filtered through a 0.22 µm filter (Amicon Ultra, 100 kDa MWCO) in order to concentrate until a final volume of ≤500 µL. Concentrated samples were loaded on qEV70 columns, previously equilibrated with filtered PBS according to the manufacturer’s protocol. Sequential fractions were collected, and uEVs-enriched fractions were pooled based on the manufacturer’s specifications.

### 2.4 Agarose beads functionalization with MSP

1.5 mL of a magnetic agarose beads suspension (Thermo Fisher Scientific, Pierce™ High-Capacity Ni-IMAC MagBeads) was incubated with a solution of His-tag Bk-MSP (20 mg/mL)^32^ in PBS (2 hours at room temperature). Upon completion, the beads were washed three times with filtered PBS and stored in PBS + 0.05% NaN_3_ until use.

### 2.5 MSP-beads-based uEVs isolation

Pre-cleared urines were processed in 20 mL batches, using 400 μL of MSP-functionalized beads. Before use, MSP-beads were aliquoted from stock and washed three times with PBS. Then, MSP-beads were vortexed for 10 seconds and incubated under mixing (inversion) for 1 hour at room temperature. Beads were then placed on a magnetic separator and washed three times with filtered PBS. EVs were eluted by incubating the beads with 400 μL of RPMI 1% DMSO 200 mM EDTA solution (“release buffer”) for 15 min at room temperature, under gentle mixing. Enriched uEVss were recovered in the supernatant after magnetic separation.

### 2.5 Transmission electron microscopy

Transmission electron microscopy (TEM) was used to examine the ultrastructure of isolated extracellular vesicles (EVs). For analysis, EV samples were properly diluted in phosphate-buffered saline (PBS) and applied to 300-mesh carbon-coated copper grids in a humidified chamber at room temperature for 15 minutes. The grids were then fixed in 2% glutaraldehyde in PBS for 10 minutes, followed by three rinses with Milli-Q water for 3 minutes each. Preparation was completed by negative staining with a 2% phosphotungstic acid solution at pH 7. Samples were observed by the transmission electron microscope Philips CM 100 TEM 80 kV; the images were captured by a Kodak digital camera.

### 2.7 Nanoparticle tracking analysis

Isolated uEVss were diluted 1:100 in a 0.1 μm filtered physiological solution and analyzed by NanoSight (Malvern Panalytical, UK) equipped with the Nanoparticle Tracking Analysis (NTA) and NTA 3.2 Analytical Software Update. A syringe pump flow of 30 was applied for each sample. Three videos of 30 s each were recorded and analyzed, yielding an estimation of the particle size distribution and concentration (particles/mL) of the EV preparations.

### 2.8 MACSPlex assay analysis

The MACSPlex exosome kit (Miltenyi Biotec, Bergisch Gladbach, Germany) was used to analyze the surface marker expression profile of MSP-isolated uEVss, as previously described^34^. As positive control, we used 250 μl the pre-cleared urine sample from which uEVss were isolated. Indeed, the MACSPlex assay can be performed both on EV preparation and on pre-cleared biofluid samples. For the assay, 1,8 x 10^9^ particles (particle number estimated considering NTA-measured particle concentration) of MSP-isolated uEVs preparation were diluted with MACSPlex buffer (MPB), reaching a final volume of 250 μl. Samples were then incubated with 15 μl of MACSPlex capture beads overnight at 4 °C. Subsequently, after washing with MPB, a mixture of APC-conjugated anti-CD9, anti-CD63, and anti-CD81 detection antibodies (provided by the kit, 5 μl of each antibody for each sample) were added to each sample. After 1 h incubation at room temperature, 500 μl of MPB were added to wash the beads, and then it was removed after one centrifugation. This step was followed by another washing with 500 μl of MPB and incubation for 15 min at room temperature. After this incubation, the samples were centrifuged and about 550 μl of supernatant were removed, leaving about 200 μl of sample in each tube. The pellet was resuspended, transferred to a flow cytometry and subjected to flow cytometric analysis using FACS Celesta (BD Biosciences, Franklin Lakes, NJ, USA). Throughout the protocol, the incubations were carried out on an orbital shaker at 450 rpm, protected from light. To wash the beads, 500 μl of MPB were added and removed after several centrifugations at 3,000 x g for 5 minutes at room temperature.

### 2.9 ELISA for Uromodulin

To evaluate the amount of uromodulin as a soluble contaminant in uEVss preparations obtained by different isolation methods (dUC, and MSP-based), as well as in unprocessed urine, a Human Uromodulin ELISA Kit (Thermo Fisher Scientific) was used. All assays were performed according to the manufacturer’s instructions, using appropriate dilutions for each sample type.

Unprocessed urine samples were diluted 1:10,000 in the dilution buffer supplied with the kit, whereas isolated uEVs preparations were diluted 1:100 in the same buffer. Samples were then processed and analyzed in parallel to allow direct comparison across conditions.

### 2.10 SiMoA assay

SiMoA assay was performed to provide high sensitivity quantification of common EV-associated markers (such as tetraspanins CD9, CD63, and CD81) in order to observe differences in terms of uEVs sub-population enrichment. Briefly, a three-step assay was set up according to the manufacturer’s instructions, as previously reported^35^. 0.1 mL of each sample were analyzed per well. Capture and detection probes consisted of a mixture of anti-tetraspanin antibodies: anti-CD9 antibody (Ancell, clone SN4/C3-3A2), anti-CD63 antibody (Ancell, clone AHN16.1/46-4-5), and anti-CD81 antibody (Ancell, clone 1.3.3.22). Results were analyzed with GraphPad Prism 6 software.

### 2.11 Fluorescence NTA

Fluorescence nanoparticle tracking analysis (F-NTA) was used to assess the co-expression of canonical EV surface markers (CD9, CD63, CD81) and to compare the co-localization profiles of uEVss isolated by the two different methods. Fluorescence signals, particle size, and concentration were measured for each sample. All uEVs samples were diluted 1:100 in Milli-Q (MQ) water before analysis. Briefly, 10 µL of uEVs suspension were incubated with fluorophore-conjugated antibodies, CD9-FITC (clone SN4 C3-3A2, Thermo Fisher Scientific), CD63-PE (clone H5C6, Thermo Fisher Scientific), and CD81-APC (clone 1D6, Thermo Fisher Scientific); at a 1:10 antibody to sample volume ratio for 1 h at room temperature. Following incubation, labeled uEVss were further diluted in MQ water to reach 1 mL, and loaded into the sample chamber of a ZetaView^®^ Quatt device (Particle Metrix, Germany), equipped with four lasers (405nm, 488nm, 520nm, 640nm) and four filters (410nm, 500nm, 550nm, 660nm) and the ZetaNavigator software 1.4.7.6. The shutter was set to 100 and the sensitivity to 70 (scatter measurement) and 90 (fluorescence measurement) for all experiments. Videos were acquired from eleven positions in the flow cell per measurement. The colocalization measurement data was based on a 60 frame video (30 frames/second) and a minimum trace length of 3 frames. Acquired videos were processed using the manufacturer’s analysis software Particle Explorer (PEX) software 4.3.4.4, applying identical settings for all samples to ensure consistency across measurements.

### 2.12 High-sensitivity flow cytometry analysis

uEVs preparations isolated with the three different methods were analyzed with Cytek Amnis CellStream (Cytek Biosciences, CA, USA) benchtop flow cytometer for the expression of CD9. For each sample, 3 x 10^8^ particles (calculated by NTA quantification) were incubated for 15 minutes protected from light with 0,75 μg/ml of Phycoerythrin (PE)-conjugated anti-CD9 antibody (Cat. No. 130-124-758, Miltenyi Biotec), reaching a total volume of 100 μl with filtered (0,1 μm pores) PBS. Subsequently, 200 μl of filtered PBS were added, and samples were vortexed for a few seconds. Parallelly, for each sample analyzed, unstained EVs, as well as EVs stained with the appropriate isotype control (Cat. No. 130-113-438, Miltenyi Biotec) at the same concentration of PE-anti-CD9 antibody. For each sample, the median fluorescence intensity (MFI) detected with the isotype control was subtracted from the MFI detected with the anti-CD9 antibody.

## 3. Results

### 3.1 MSP-based isolation retains efficient yields and increases uEVs purity compared to dUC

Magnetic (agarose) beads were selected as the solid support for MSP conjugation in this study because they enable rapid, instrument-free separation with minimal sample handling. Bk-MSP peptide was selected as previously validated in other settings, including cell-conditioned media and plasma^31,32^.

We investigated uEVs isolation using urine samples collected from five healthy donors, directly compared MSP-based isolation with dUC, used as the reference method. To exclude potential biases associated with differences in resuspension or elution buffers, dUC-isolated uEVss were analyzed in parallel after resuspension in either the standard dUC buffer or the MSP release buffer. As comparable results were obtained across all characterization assays (Figure S1), subsequent analyses are presented using data from uEVss resuspended in the standard dUC buffer.

The resulting uEVs preparations were first evaluated in terms of their physical properties. Nanoparticle tracking analysis (NTA) revealed comparable size distributions across both isolation methods, with similar mean (∼150 nm) and mode (∼100 nm) diameters (Fig. 2B–D). NTA-based particle quantification showed a modest reduction in particle counts in MSP-isolated samples compared with dUC preparations (Fig. 2A). However, it is well known that non-EV particles or protein aggregates can contribute to counting events in NTA analysis; this difference can likely be ascribed to a higher level of co-isolated contaminants in dUC preparations rather than reduced EV recovery by MSP-based isolation ^36–38^. Transmission electron microscopy (TEM) further confirmed the presence of intact vesicles following release from MSP-functionalized beads, displaying the characteristic cup-shaped morphology typical of EVs and showing minimal evidence of contaminating material (Fig. 2E).

**Figure 2:**
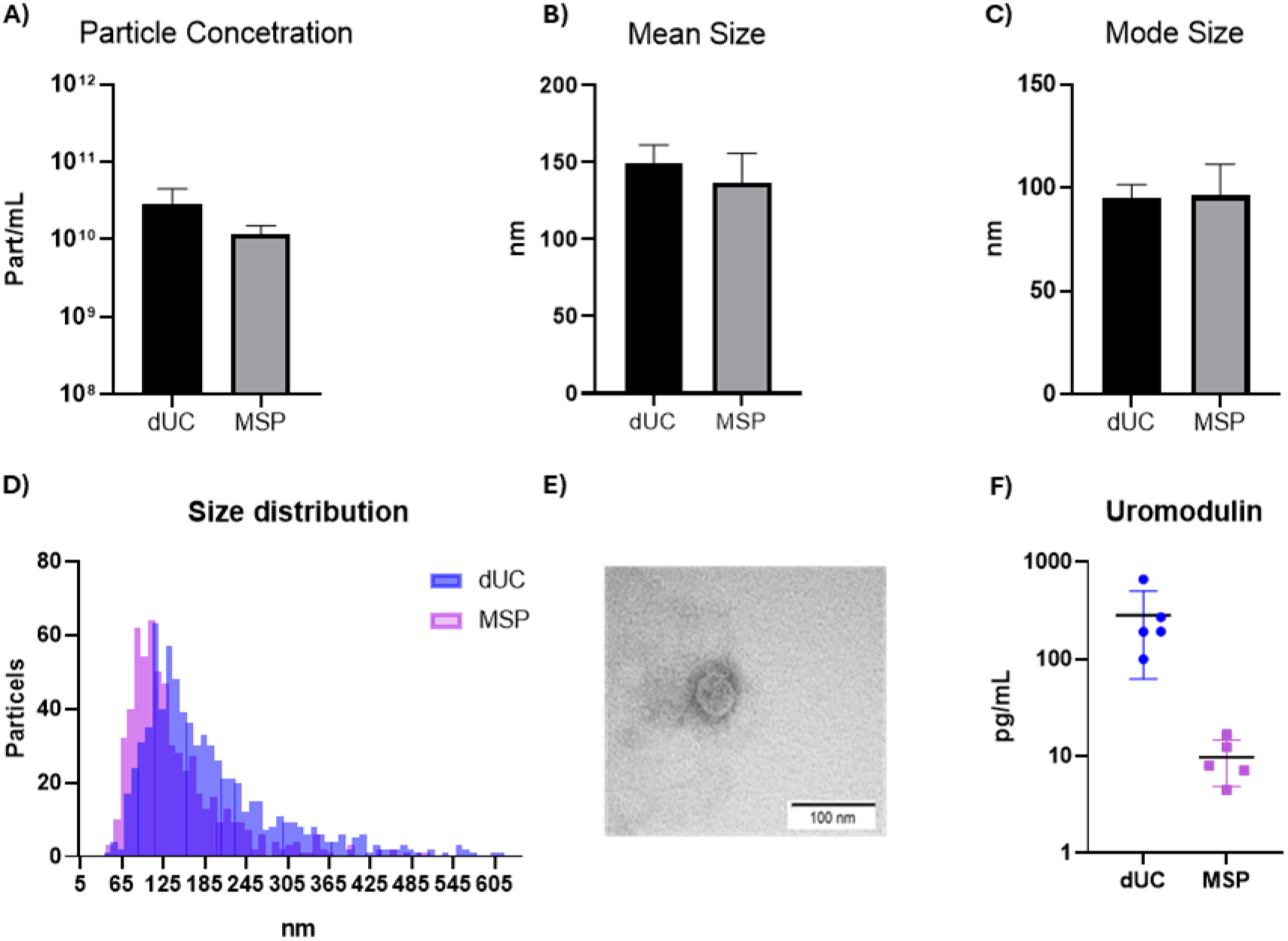
A) Particle concentration estimated by NTA of dUC-isolated uEVss and MSP-isolated uEVss (N=5). B&C) Mean and mode size estimated by NTA of dUC-isolated uEVss and MSP-isolated uEVss (N=5); D) Representative size distribution profile of dUC-isolated uEVss and MSP-isolated uEVss; E) Representative picture of TEM of isolated MSP-isolated EVs (scale bar: 100 nm); F) Uromodulin concentration measured by ELISA in dUC-isolated uEVss and MSP-isolated uEVss (N=5).

To more precise assessment of uEVs purity, we quantified the amount of residual uromodulin (THP) by using a commercial ELISA test (see protocols section), the most common contaminant protein in uEVs preparations. Reported results in Figure 2F clearly show that MSP-based EV isolation was more effective in reducing uromodulin residual amount, with a nominal >30 fold purity increase (average of THP after dUC ≈299 pg/mL vs. ≈10 pg/mL after MSP isolation) over dUC (Figure S2). Based on the combined data from size-distribution analysis, particle counting, and uromodulin quantification, the data suggest that MSP-based uEVs isolation provides similar recovery and enhanced purity compared with dUC—at least with respect to uromodulin content.

### 3.2 MSP-based isolation yields an uEVs tetraspanin profile comparable to standard methods

Following the physical characterization of the isolated uEVs preparations, we next evaluated their molecular profile by assessing the expression of established EV surface markers. As an initial validation of MSP-based uEVs isolation, a multiplexed bead-based assay (MACSPlex) was employed to profile 37 EV-associated surface markers and assess whether MSP capture preserves the molecular representativeness of uEVss relative to the corresponding pre-cleared urine samples. MSP-isolated uEVss exhibited robust expression of the canonical EV tetraspanins CD9, CD63, and CD81, together with established uEVs-associated markers, including CD133-1, CD24, and CD326 (Fig. 3A). Importantly, the overall marker profile closely mirrored that of the corresponding pre-cleared urine used as positive control, indicating that MSP-based isolation does not selectively enrich or deplete specific EV subpopulations (Fig. 3A).

**Figure 3:**
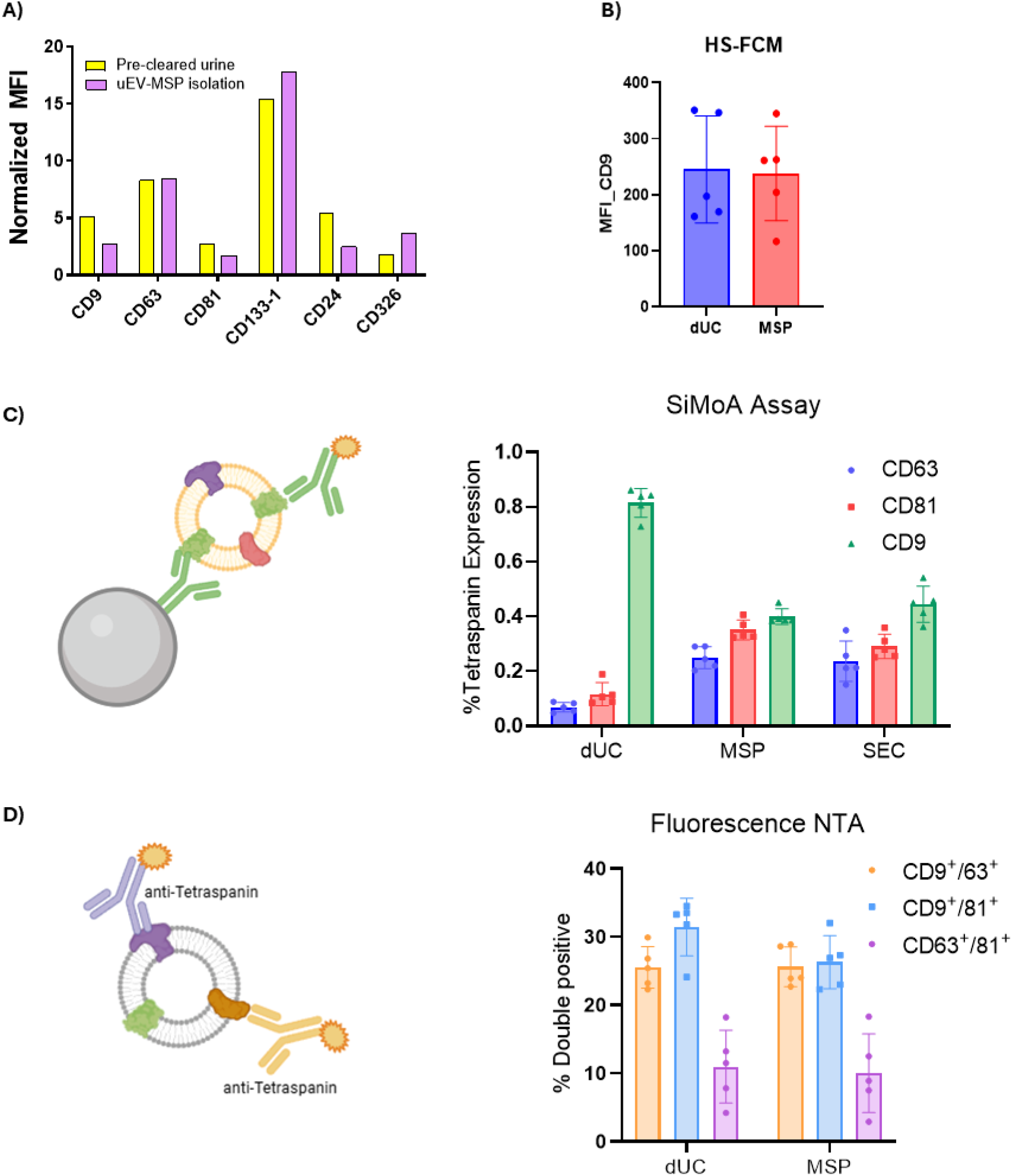
Comparative analysis of tetraspanin expression and co-localization in isolated uEVss. A) MACSPlex analysis in pre-cleared urine samples (yellow bars) and in MSP-isolated uEVss (purple bars). Data were normalized to the median fluorescence intensity of all detectable markers; B) CD9 Median Fluorescence Intensity in the small EV isolated measured using HS-FCM technology comparing dUC and MSP; C) Schematic representation of the single-tetraspanin SiMoA assay; corresponding plots show the average enzyme per bead (AEB) signal for individual tetraspanins (CD9, CD63, and CD81) in uEVss isolated by dUC, SEC, and MSP-functionalized beads (N = 5). B) Sketch of EV-labeling with mAbs anti-tetraspanin for F-NTA analysis; percentage of events that colocalized for two different tetraspanin simultaneously.

Focusing on the canonical CD9 as urinary EV marker, high-sensitivity flow cytometry (HS-FCM) was performed (Fig. 3B). Unlike MACSPlex analysis, which adopts bulk bead-based immunocapture, HS-FCM results are filtered through optimized trigger and scatter settings to selectively quantify fluorescence events associated with nanosized particles. When an appropriate size gate was applied, no significant differences in the proportion of CD9-positive events were detected between dUC- and MSP-isolated uEVss (Fig. 3B).

Subsequently, we comprehensively compared the preservation of molecular surface marker fingerprints in isolated uEVss, focusing on the canonical tetraspanins (CD9, CD63, CD81). To this end, we employed advanced analytical platforms, including single molecule array analysis (SiMoA^®^)—an ultrasensitive bead-based digital ELISA—and fluorescence NTA, which provides single-particle–level information. For this purpose, uEVss from five healthy donors were isolated as described in the Methods section, and their tetraspanin expression profiles were evaluated under both dUC and MSP isolation conditions.

For SiMoA analysis, assays were configured using the same antibody clone for both EV capture and detection, enabling the quantification of the relative abundance of individual tetraspanins within each sample (Fig. 3C).

Overall, the tetraspanin abundance profiles obtained with dUC and MSP-based isolation showed highly comparable trends, indicating preservation of the native EV marker distribution across the two methods. Notably, dUC samples displayed a relatively increased CD9 signal compared with MSP-isolated uEVss. To further investigate this observation, size-exclusion chromatography (SEC) was included as an additional isolation strategy for the same set of urine samples. SEC represents a distinct EV separation principle, allowing us to benchmark the results against an orthogonal and widely used approach. MSP and SEC exhibited closely comparable tetraspanin profiles, and overall, SiMoA analysis demonstrated consistent results across all three isolation methods, indicating that CD9-positive EVs constituted the predominant fraction (Fig. 3C). These findings suggest that MSP-based isolation preserves the relative abundance of canonical EV markers without introducing substantial subpopulation bias.

Then, we further expanded uEVs characterization at the single-particle level by using Fluorescence NTA (F-NTA) to quantify vesicles simultaneously positive for pairs of tetraspanin markers (Fig 3D). Specifically, we measured the proportion of CD9/CD63, CD9/CD81, and CD63/CD81 double-positive vesicles across uEVss isolated by dUC and by MSP-functionalized beads. The analysis showed that, apparently, the fraction of double-positive vesicles was conserved across both isolation methods, with no evidence of selective enrichment or depletion of any tetraspanin co-expressing subpopulations. Additionally, the overall number of vesicles positive for at least two tetraspanins remained comparable between MSP-functionalized beads and dUC-isolated samples derived from the same urine source (Figure S3). Overall, within the uEVs subsets falling within the overlapping working range of both technologies, some inter-sample variability was observed. However, this variability was comparable across the two isolation methods and is more likely to reflect biological differences in tetraspanin expression among donors rather than methodological effects.

## 4. Discussion

In this study we examined MSP-based affinity capture for the isolation of urinary EVs, focusing on its feasibility as an alternative to conventional methods and, critically, on its potential to overcome key limitations that currently restrict scalable, reproducible, and clinically translatable uEVs isolation workflows.

When benchmarked against differential ultracentrifugation (dUC)—still regarded as a reference method in many EV workflows^39–42^—MSP-functionalized beads provided uEVss with highly comparable size distribution, recovery yield, and general EV-associated markers profile. At the same time, MSP-based enrichment markedly reduced uromodulin contamination, addressing a major confounding factor in uEVs research. Given the recognized tendency of uromodulin to co-pellet or co-precipitate with EVs^43,44^, the ability of MSP capture to selectively retain vesicles while minimizing non-vesicular proteins may enhance the reproducibility and downstream performance of uEVs preparations.

An additional advantage of this approach is that, unlike several commercially available bead-based isolation methods, the beads can be detached and separated from the isolated uEVss. This feature may help overcome a common limitation of bead-based approaches, in which EVs remain associated with the beads, thereby constraining downstream analytical of functional applications. Moreover, these latter approaches are usually based on an immune-capture, which depends on marker expression. At variance, the MSP method does not introduce detectable bias toward or against specific tetraspanin-defined EV subsets. Furthermore, it yields a surface-marker profile highly comparable to SEC—another widely accepted reference method in uEVs isolation, often associated with high purity. Likewise, no evident bias emerged when benchmarking MSP results against dUC within the analytical window defined by the operational range of common nanoanalytical technologies. These observations were consistently supported by orthogonal analyses, including SiMoA, fluorescence-NTA, and high-sensitivity flow cytometry. Notably, SiMoA analysis revealed an increased signal for CD9 in dUC samples when evaluating single-marker abundance, suggesting a relative enrichment of CD9^+^ vesicles. However, this apparent difference was not maintained when CD9 expression was assessed in combination with other EV canonical markers, such as CD63 and CD81. Indeed, F-NTA analysis, which enables the detection of vesicles co-expressing markers, showed comparable levels of double-positive particles across isolation methods. Collectively, our data support MSP-based affinity as an efficient strategy for isolating urinary extracellular vesicles, preserving the native heterogeneity of uEVss without introducing selective enrichment.

Beyond this early analytical assessment, examining additional operational parameters further highlights the potential value of MSP-based isolation (Figure 1). Total process time for MSP isolation (uEVs capture, washes, and elution) is approximately 1.5–2 hours, compared with ≥4–5 hours for dUC (including long centrifugation cycles and pellet handling). Estimated hands-on time is reduced to 10–15 minutes per batch, compared with the 45–60 minutes typically required for dUC, which includes sample balancing, tube preparation, pellet recovery, and rotor maintenance. Moreover, MSP-based isolation relies on simple bead mixing and magnetic separation using standard wet-lab equipment, without the need for costly or specialized instrumentation as the dUC. Bead volumes can be proportionally adjusted to accommodate higher or lower sample inputs while maintaining identical workflow steps, enabling flexible scalability. This feature is expected to minimize operator-to-operator and sample-to-sample variability—an intrinsic limitation of pellet-based ultracentrifugation workflows. Overall, these characteristics point toward improved process robustness, an increasingly relevant requirement in the context of EV workflow standardization.

We herein described and validated a method for uEVs isolation which is of substantial relevance for therapeutic applications and biomarker discovery. Moreover, the characterization of the uEVss captured by the MSP could represent a starting point for the development of MSP-based, isolation-free EV analytical platforms. Ideally, MSP-driven capture could potentially replace the immune-capture currently employed in several analytical techniques, thereby avoiding the biased isolation of antigen-expressing EV subpopulations. In this context, such strategies may facilitate the integration of EV analysis into diagnostic and liquid biopsy applications.

## 5. Conclusion

Collectively, the results of this study demonstrate that MSP-based isolation enables the recovery of uEVs with preserved molecular and biophysical characteristics while significantly reducing co-isolated urinary contaminants, particularly uromodulin, compared with dUC. Orthogonal characterization approaches consistently showed that MSP capture preserves the native distribution of canonical EV markers without introducing detectable subpopulation bias.

In addition to improved purity, the MSP workflow offers practical advantages in terms of simplicity, scalability, and compatibility with standard laboratory infrastructure, avoiding the lengthy processing times and specialized instrumentation required by dUC. Taken together, these findings support MSP-based affinity capture as a robust and clinically relevant strategy for standardized uEV isolation and downstream biomarker analyses.

## Supporting information

Supplementary Information of the main manuscript

## In memory of Marina

This research was partially funded by the European Union through Horizon 2020 research and innovation program under grant agreement No. 951768 (project MARVEL). Work was also supported by PNRR MUR – M4C2, CN3 National Center for Gene Therapy and Drugs based on RNA Technology - Spoke 8, MUR CN00000041 CUP D13C22001310001.

