## Supplementary Information of the main manuscript for "Enhanced Workflow for Urinary Extracellular Vesicle Isolation Using Membrane-Sensing Peptides"

**Supplementary Figure 1:**

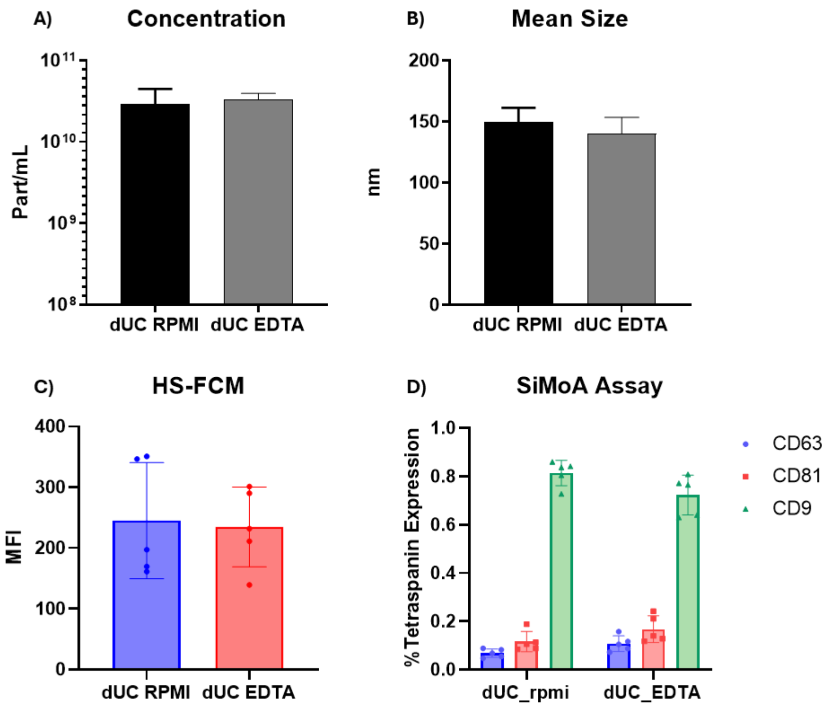


**Assessment of potential effects of buffers on uEVs characterization.**

To evaluate whether buffer composition introduces variability in isolated uEVs, a side-by-side comparison was performed using dUC-isolated samples resuspended either in standard RPMI buffer or in the MSP release buffer.

A&B) Particle concentration and mean size measured by measured NTA for dUC-isolated uEVs resuspended in the two different buffers (N = 5). C) Median fluorescence intensity of CD9 measured by high-sensitivity flow cytometry. D) Pan-tetraspanin SiMoA assay showing the average enzyme per bead (AEB) signal for CD9, CD63, and CD81. Across all analyses, comparable results were observed between the two conditions, indicating that buffer composition does not significantly affect uEVs characterization.

**Supplementary Figure 2:**

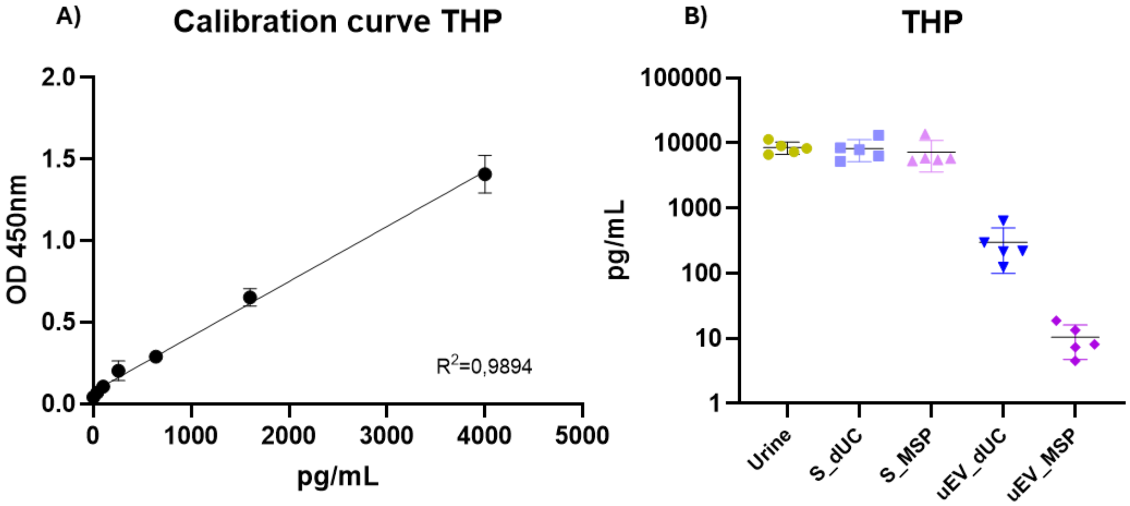


**Quantification of Uromodulin across uEVs isolation workflows.**

THP levels were measured to assess co-isolated contaminants in both, dUC and MSP-based preparations.

A) ELISA calibration curve used to determine THP concentration (pg/mL). B) THP levels measured in five samples across different stages of the isolation process, including starting urine, supernatant (S) the uEVs-depleted fraction and, final uEVs preparations. Lower THP levels in MSP-isolated samples indicate improved removal of soluble contaminants.

**Supplementary Figure 3:**

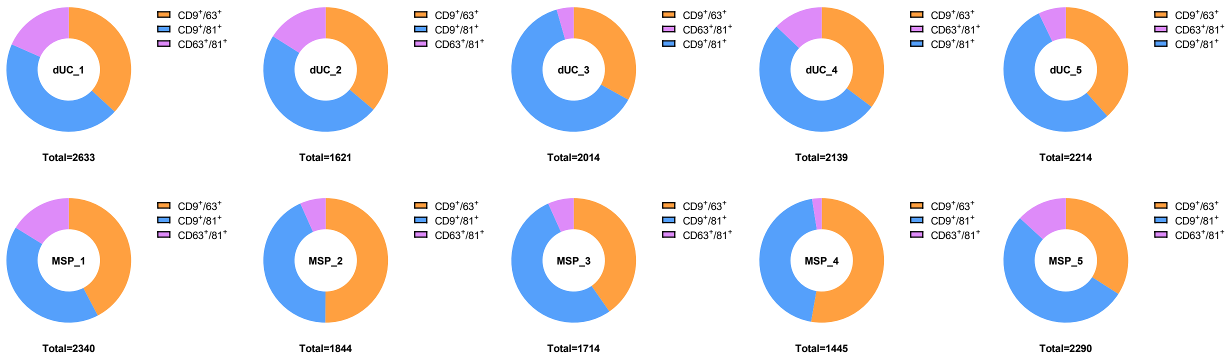


**Comparison of double-tetraspanin positive uEVs across isolation methods.**

The total number of vesicles co-expressing at least two tetraspanins was comparable between dUC and MSP-functionalized bead isolated samples derived from the same urine source. Moreover, donut plot representations further illustrate that the relative distribution of double-positive vesicle populations is preserved across the two isolation methods for each individual sample.
